# Common micro- and macroscale principles of connectivity in the human brain

**DOI:** 10.1101/2021.09.14.459604

**Authors:** Lianne H. Scholtens, Rory Pijnenburg, Siemon C. de Lange, Inge Huitinga, Martijn P. van den Heuvel, Netherlands Brain Bank (NBB)

## Abstract

The brain requires efficient information transfer between neurons and between large-scale brain regions. Brain connectivity follows predictable organizational principles: at the cellular level, larger supragranular pyramidal neurons have larger dendritic trees, more synapses, more complex branching and perform more complex neuronal computations; at the macro-scale, region-to-region connections are suggested to display a diverse architecture with highly connected hub-areas facilitating complex information integration and computation. Here, we explore the hypothesis that the branching structure of large-scale region-to-region connectivity follows similar organizational principles as known for the neuronal scale. We examine microscale connectivity of basal dendritic trees of supragranular pyramidal neurons (300+) across ten cortical areas in five human donor brains (1M/4F). Dendritic complexity was quantified as the number of branch points, tree length, spine count, spine density and overall branching complexity. High-resolution diffusion-weighted MRI was used to construct ‘white matter trees’ of cortico-cortical wiring. Examining the complexity of the resulting white matter trees using the same measures as for dendritic trees shows multimodal association areas to have larger, more complexly branched white matter trees than primary areas (all p<0.0001) and regional macroscale complexity to run in parallel with microscale measures, in terms of number of inputs (r=0.677, p=0.032), branch points (r=0.790, p=0.006), total tree length (r=0.664, p=0.036) and branching complexity (r=0.724, p=0.018). Our findings support the integrative theory that brain connectivity is structured following similar ‘principles of connectivity’ at the neuronal and macroscale level, and provide a framework to study connectivity changes in brain conditions at multiple levels of brain organization.

## Introduction

Efficient transfer of information between neurons and regions of the human brain is essential for perception, cognition and behavior. Understanding how brain wiring on both cellular and regional scales is shaped to facilitate such efficient information processing is a central topic in the field of neuroscience.

Variation in neuronal connectivity is widely believed to be a key factor underlying regional differences in information processing (Schüz and Miller, 2003) and regional functional specialization (Amunts and Zilles, 2015). A region’s role and capacity for information processing is suggested to be an important mix of cortical type (Barbas and Rempel-Clower, 1997), neurotransmitter receptor densities (Zilles and Palomero-Gallagher, 2017), intracortical myelination (García-Cabezas et al., 2017), cortical and glial density (Collins, 2011) and dendritic complexity of pyramidal neurons (Elston, 2003b). Pyramidal neurons in particular are an important class of neurons in the context of neural integration and cortico-cortical connectivity (Schüz and Miller, 2003). Larger, more complex layer II/III pyramidal neurons are thought to display a higher capacity for information integration, and are typically found in regions involved in multimodal cognitive processing (Jacobs et al., 2001; Elston, 2003a). Pyramidal neuron connectivity and dendritic branching result from a balance between space and energy cost (Wen and Chklovskii, 2008; Wen et al., 2009; Cuntz et al., 2010; Schröter et al., 2017) and maximization of the neuron’s connectivity repertoire (Wen et al., 2009). Importantly, the axonal projections of supragranular pyramidal neurons comprise a large portion of the white matter bundles interconnecting large-scale brain regions (Jacobs et al., 1997; Jacobs et al., 2001; Kaas et al., 2002; Barbas, 2015), forming the macroscale wiring network of the human brain (Bullmore and Sporns, 2009; Sporns, 2009; van den Heuvel and Sporns, 2011). Pyramidal neurons thus form an important class of neurons operating at both the microscale and macroscale level of brain connectivity.

MRI studies in turn have shown brain connectivity and macroscale functional networks to display distinct organizational principles with short communication relays (Bullmore and Sporns, 2009), structural and functional community structure (Meunier et al., 2010; Yeo et al., 2011) and the formation of central ‘hubs’ for topological global integration of information (Tomasi and Volkow, 2011; van den Heuvel and Sporns, 2011). These network properties too are suggested to be shaped by a trade-off between ‘cost’ and ‘efficiency’ (Bassett et al., 2009; Bullmore and Sporns, 2012; van den Heuvel et al., 2012), observations contributing to a central question in the field of how different scales of brain connectivity relate to each other (Perin et al., 2011; Van Wedeen et al., 2012; Barbas, 2015; van den Heuvel and Yeo, 2017; Scholtens and van den Heuvel, 2018; García-Cabezas et al., 2019; Goulas et al., 2019).

We examined the relationship between these two levels of brain organization with the goal to examine whether and if so how the organizational principles that govern neuronal branching and complexity also shape the macroscale region-to-region connectivity. Across ten cortical regions, we reconstructed three-dimensional morphology of microscale dendritic branching and macroscale white matter wiring (Elston and Rosa, 1997; Jacobs et al., 1997; Elston et al., 1999; Jacobs et al., 2001). We show that macroscale connectivity is organized according to principles similar to those of microscale neuronal connectivity, with larger, more branched connectivity in association areas. In a direct comparison, our findings show that micro and macroscale branching complexity strongly go hand in hand across brain areas, supporting the hypothesis of shared principles of connectivity across scales.

## Methods

### Human post-mortem tissue

Cortical brain samples of five human donors were collected from the Netherlands Brain Bank (NBB, Amsterdam, the Netherlands; www.brainbank.nl)(See for demographics of the donors, Table 1 & 2). Informed consent to perform autopsies and for the use of tissue and clinical data for research purposes was obtained by the NBB from donors and approved by the Ethical Committee of the VU University medical center (VUmc, Amsterdam, The Netherlands). A total of 42 post-mortem samples was obtained across 10 cortical regions: gyrus frontalis superior, gyrus frontalis medius, gyrus cingularis anterior, precuneus, gyrus insularis brevis, gyrus precentralis, gyrus postcentralis, gyrus occipitalis superior, occipital pole, lateral gyrus parietalis superior (Figure 1A)(Table 2), covering multimodal association areas, unimodal, primary areas and limbic cortex.

**Table 1.**
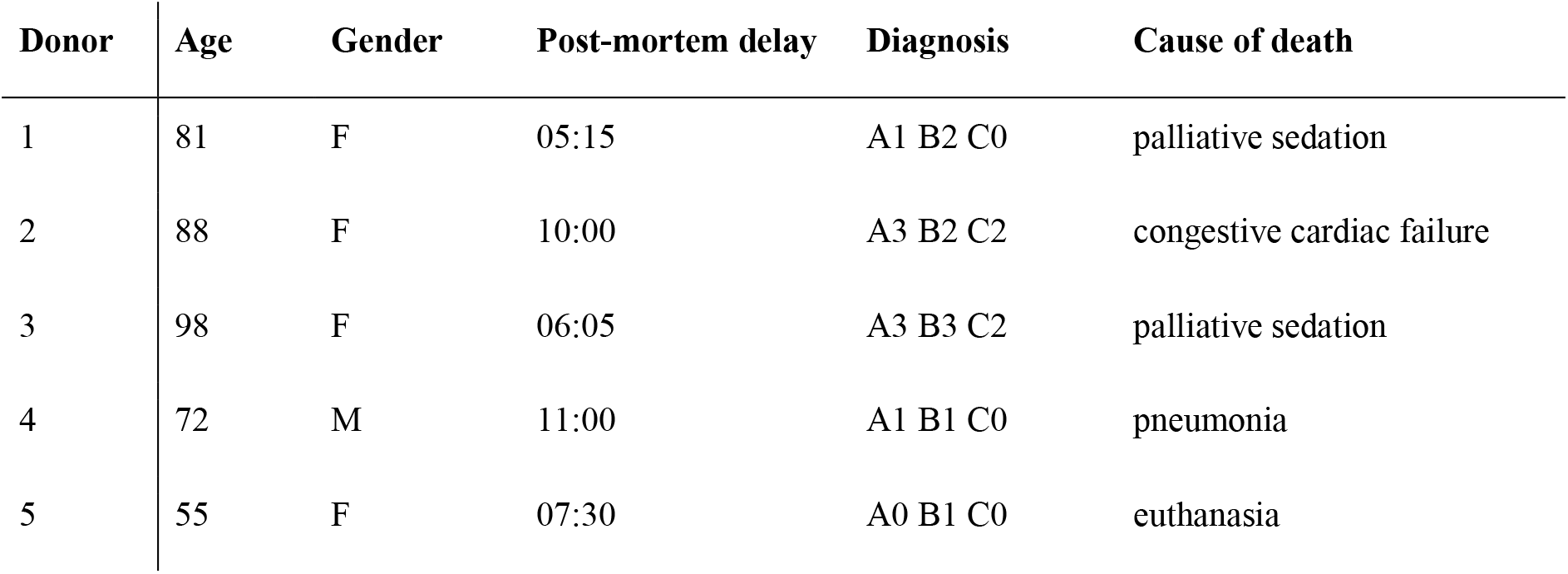
Donor demographics. The table describes for each donor (from left to right), the age at death, gender, post-mortem delay (hours:minutes), ABC neuropathological score (describing the severity of amyloid-β plaques (A), Braak stage (B) and neuritic plaques (C)), as well as the reported cause of death.

| Donor | Age | Gender | Post-mortem delay | Diagnosis | Cause of death |
| --- | --- | --- | --- | --- | --- |
| 1 | 81 | F | 05:15 | A1 B2 C0 | palliative sedation |
| 2 | 88 | F | 10:00 | A3 B2 C2 | congestive cardiac failure |
| 3 | 98 | F | 06:05 | A3 B3 C2 | palliative sedation |
| 4 | 72 | M | 11:00 | A1 B1 C0 | pneumonia |
| 5 | 55 | F | 07:30 | A0 B1 C0 | euthanasia |

**Table 2.**
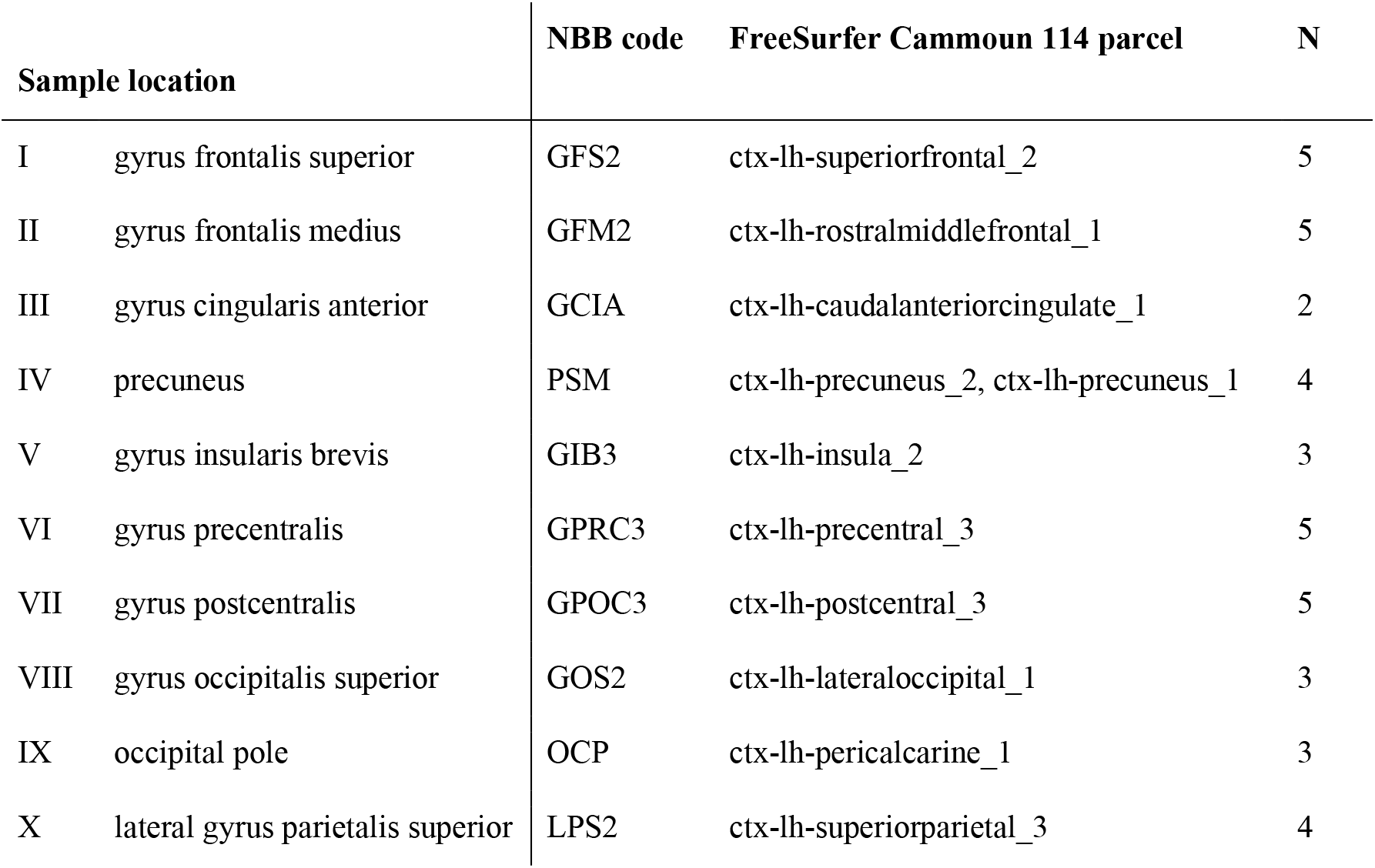
Included tissue samples. The table lists the ten included sample locations, all resected from the left hemisphere, describing (from left to right) the sample number, textual description of the sample location, the corresponding Netherlands Brain Bank (NBB) sample code, the mapped Cammoun 114 FreeSurfer parcel(s) and the number of donors the sample was successfully obtained from.

| Sample location |  | NBB code | FreeSurfer Cammoun 114 parcel | N |
| --- | --- | --- | --- | --- |
| I | gyrus frontalis superior | GFS2 | ctx-lh-superiorfrontal_2 | 5 |
| II | gyrus frontalis medius | GFM2 | ctx-lh-rostralmiddlefrontal_1 | 5 |
| III | gyrus cingularis anterior | GCI A | ctx-lh-caudalanteriorcingulate_1 | 2 |
| IV | precuneus | PSM | ctx-lh-precuneus_2, ctx-lh-precuneus_1 | 4 |
| V | gyrus insularis brevis | GIB3 | ctx-lh-insula_2 | 3 |
| VI | gyrus precentralis | GPRC3 | ctx-lh-precentral_3 | 5 |
| VII | gyrus postcentralis | GPOC3 | ctx-lh-postcentral_3 | 5 |
| VIII | gyrus occipitalis superior | GOS2 | ctx-lh-lateraloccipital_1 | 3 |
| IX | occipital pole | OCP | ctx-lh-pericalcarine_1 | 3 |
| X | lateral gyrus parietalis superior | LPS2 | ctx-lh-superiorparietal_3 | 4 |

**Figure 1.**
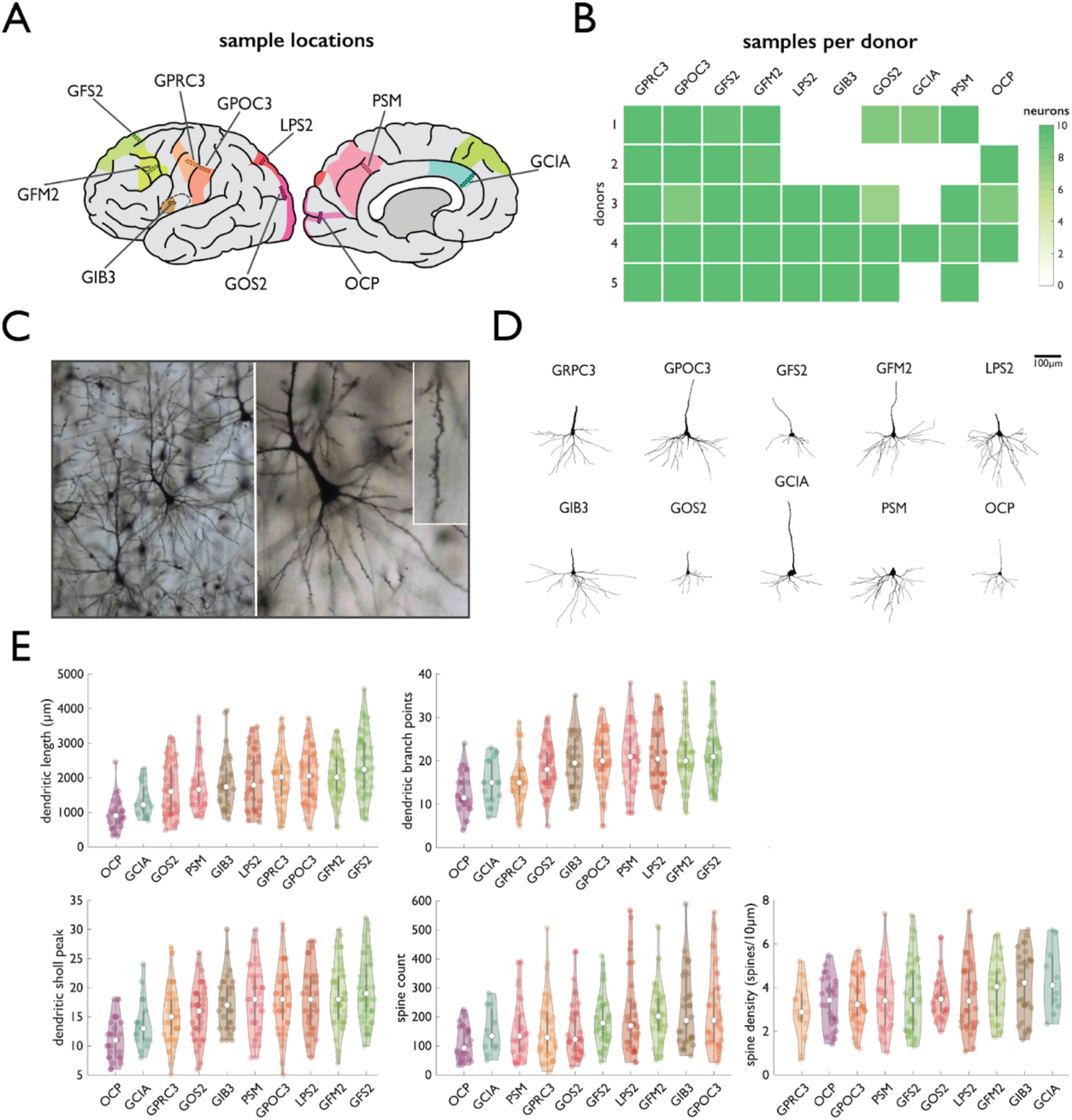
Regional variation in microscale basal dendritic tree organization. Panel **A** shows a schematic depiction of the locations of the ten included cortical samples: GPRC3 - gyrus precentralis, GPOC3 - gyrus postcentralis, GFS2 - gyrus frontalis superior, GFM2 - gyrus frontalis medius, LPS2 - lateral gyrus parietalis superior, GIB3 - gyrus insularis brevis, GOS2 - gyrus occipitalis superior, GCIA - gyrus cingularis anterior, PSM - precuneus, OCP - occipital pole. Panel **B** depicts the number of reconstructed neurons in each tissue sample, for each of the five donor brains. White squares indicate samples not obtained from that donor, darker shades of green indicate a larger number of neurons included in a sample (max 10 per sample). **C** Shows a representative example of Golgi-Cox stained layer III pyramidal neuron imaged at 10x (left) 20x (middle) and 40x (right). Panel **D** shows an example neuron reconstruction for each of the included brain regions (numbered to match panel A) of donor 4. Panel **E** depicts the region ranking in terms of the four measures of wiring complexity, from left to right, top to bottom: basal dendritic length (in μm), dendritic branch points, peak dendritic Sholl intersections, spine count and spine density (in spines per 10μm).

### Tissue processing

Tissue samples were dissected in 0.5 cm thin blocks perpendicular to the gyral surface spanning the width of the gyrus of interest (Figure 1A), while avoiding any unnecessary pressure on the tissue to minimize staining artifacts later in the Golgi-Cox (Cox, 1891; Ramon-Moliner, 1970) staining process. Brain tissue was moved directly into light-proof jars containing Golgi-Cox solution, for simultaneous fixation and impregnation (see Supplemental Materials and (Glaser and Van der Loos, 1981) for full protocol). After 2.5 - 3 weeks, test sections of each tissue sample were processed to assess the progression and quality of the Golgi-Cox impregnation. In 39 of the 42 tissue samples the full extent of dendritic branches was stained and spines were visible on distal branches, the remaining three samples showed substantial staining artefacts and could not be used for neuron reconstruction. The remaining 39 well-stained samples (Figure 1B) were removed from the Golgi-Cox solution, rinsed and taken to the next processing steps, dehydrating and celloidin embedding of the tissue in preparation for sectioning. The resulting tissue blocks were cut in 180 μm sections, cut perpendicular to the cortical surface using a sledge microtome (Leica / Reichert-Jung Polycut S) and transferred to ethanol 70% in individual compartments of a Teflon disk for the free-floating development and dehydration process.

### Cell selection and quantification of neuronal complexity

#### Localization

Every 6th section of each tissue block was Nissl stained in order to provide a staining of all neuronal cell bodies in the section. Using these Golgi-Nissl double stained sections the lower boundary of layer III was marked. Neurons in adjacent Golgi-stained sections were selected for morphological reconstruction using the marked lower layer III boundary for reference, selecting neurons at a cortical depth just above the boundary line.

#### Neuron morphology

Pyramidal cells from each sample were selected from the set of neurons with their soma located in supragranular layer III, as defined in the nearest Nissl counter-stained section. Pyramidal neurons were included when *i*. the soma was located in the center of the tissue section; *ii.* at least two basal dendrites could be observed that each branched at least twice; *iii.* there were no broken branches; *iv.* no branches were occluded by staining artifacts or other cells. A 3D reconstruction of the neurons was made using Neurolucida (version 11, MicroBrightfield, Williston), using a 40x oil-immersion objective (Carl Zeiss Axioskop microscope). Pyramidal cells were manually traced using the neuron tracing tools included in Neurolucida, automatically registering the dimensions and branch order of each reconstructed neuron part. Tracing began with the unmyelinated initial segment of the axon as a landmark point for placement of the traced reconstruction, followed by an outline of the soma of the neuron, after which the main branch of the apical dendrite was reconstructed for orientation purposes. Finally, the basal dendrites including all branches were traced. Neuron reconstruction was performed by trained experts LHS and RP at the Netherlands Brain Bank and resulted in on average 75.2 neurons per donor (ranging from 49 to 99) and 37.6 per included brain region (ranging from 18 to 50), yielding a total of 376 reconstructed pyramidal neurons.

#### Spine count and spine density

The quality of the Golgi-Cox impregnation was assessed for each sample, evaluating the extent to which spines were fully impregnated, evidence of over-impregnation that could impede accurate quantification of spines, and/or staining artifacts occluding spines. Samples of the first included donor (donor 1) were used for optimization of the spine quantification protocol and not further included in the analysis. For the samples of the remaining four donors, where quality of staining was scored to be sufficient (91% of samples), the best visible, longest basal dendrite of each included neuron was selected for spine quantification, resulting in a total of 281 neurons with spine quantification included. Spines were marked when *i.* they had a visible head; and *ii.* were visibly attached to the dendrite.

### Analysis of neuronal morphology

Information on dendritic branching was extracted, and Sholl analysis was performed on the reconstructed neuron morphology using the tools natively included in Neurolucida. The following representative measures were selected based on previous studies (see e.g. (Scholtens et al., 2014) and Supplemental Figure S1), including *i.* total basal dendritic length; *ii.* number of branch points; *iii.* Sholl peak branching complexity; *iv.* spine count; *v.* spine density. Spine density was computed by dividing the number of counted spines by the length (in μm) of the dendrite on which the spines were quantified and multiplying with ten to obtain spine density per 10μm. Neuron reconstructions will be made available to the NeuroMorpho.org repository of digitally reconstructed neurons.

### Macroscale connectivity

#### MRI preprocessing and connectome reconstruction

High-resolution diffusion-weighted scans of 489 subjects of the Human Connectome Project (HCP; (Van Essen et al., 2013); Q3 Subjects Release; age 22-35 years, male and female combined) were used to reconstruct maps of the macroscale human connectome, mapping white matter pathways between cortical areas (de Reus and van den Heuvel, 2014). DWI parameters included 1.25 mm isotropic voxel size, TR/TE 5520/89.5 ms, 270 diffusion directions with diffusion weighting 1000, 2000, or 3000 s/mm^2^ (Van Essen et al., 2013). For each individual, preprocessing of the diffusion-weighted images included realignment, eddy current and susceptibility distortion correction, followed by a voxel-wise reconstruction of diffusion profiles using generalized q-sampling imaging and whole-brain streamline tractography using CATO (de Lange and van den Heuvel, 2021). Fiber tracing started from all white matter voxels with 8 seeds per voxel, with fiber tracing following the main diffusion direction of each voxel until reaching one of the stopping criteria (i.e. the fiber was about to exit the brain mask, made a turn of > 45degrees and/or reached a voxel with low fractional anisotropy < 0.1). High-resolution anatomical T1 scans were used for cortical segmentation and FreeSurfer (Fischl et al., 2004) parcellation of the cortical sheet of each individual subject into 114 distinct regions (57 regions per hemisphere) using the Cammoun 114 subdivision of the Desikan-Killiany atlas (Desikan et al., 2006; Cammoun et al., 2012). The individual cortical parcellation was overlaid with the subject’s whole-brain tractography to form a 114 × 114 connectivity matrix, describing all pairs of cortical regions and their reconstructed pathways (van den Heuvel et al., 2019). To match the microscale neuronal reconstructions sampled exclusively from the left hemisphere, connections of the left hemisphere were taken for further analysis, resulting in a 57 × 57 connectivity matrix.

### White matter branch reconstruction

To examine parallels between the branching structure of pyramidal neurons and the branching structure of the white matter fibers we introduced the same format to describe neuron morphology to now describe the organization of reconstructed ‘white matter trees’. We developed the following approach.

#### Step 1

For each of the 57 left hemisphere regions, a subject’s whole-brain tractography was overlaid with the individual regional parcellation to select all reconstructed streamlines for that region.

#### Step 2

The trajectory of each streamline of a region was followed in steps of 5 mm to determine where streamlines could be grouped into a white matter branch, drawing a sphere of radius 5 mm at each step. Neighboring streamlines were bundled together in locations where the radii of two spheres intersected and the angle between two potentially neighboring fibers was smaller than 90°, indicating that they projected in the same direction. To account for streamlines that follow a more winding trajectory than others and thus traverse a smaller distance in a given number of steps, spheres were allowed to intersect with neighboring spheres of equal or lower step count. Branches consisting of more than three fibers were included for further processing, computing their branch diameter and coordinates.

#### Step 3

All reconstructed branches of a region together were used to construct a ‘white matter tree’ of that cortical area. The center of the cortical area was then placed at the zero-coordinate point of the white matter tree. Each white matter branch was described at intervals of 5 mm, listing at each point the coordinates relative to the central point, together with the branch diameter and the previous linked point along the branch. The resulting white matter tree was saved in the widely used *SWC* format for neuron morphologies.

#### Step 4

Per area, the branches of the connectivity tree were examined for their level of complexity, an analysis parallel to the branching complexity analyses performed to determine connectivity complexity at the neuronal scale. White matter trees were analyzed using the TREES toolbox for quantitative analysis (Cuntz et al., 2010), computing a white matter tree equivalent of ‘total length’, ‘number of branch points’ and ‘peak branching complexity’. Total length was taken as the sum of the length of all branches of the white matter tree (in mm). The number of branch points represents the number of locations where previously parallel streamlines diverge. Peak branching complexity was computed as the maximum number of intersections of the white matter tree’s branches with concentric circles around the tree’s center point (cortical parcel).

## Cross-scale analyses

Post-mortem cortical sample locations were manually mapped to FreeSurfer’s fsaverage brain to determine corresponding macroscale Cammoun 114 FreeSurfer areas (Figure 1A, Table 2)(Fischl, 2012). The spatial location of the post-mortem tissue samples was mapped to their corresponding cortical region, matching the Cammoun-114 atlas (Figure 2). For the precuneus tissue samples (see Figure 1, Table 2) the resection as shown on the autopsy photographs was located on or close to the border between two cortical parcels. These samples were assigned to both precuneus sub-parcels, and their macroscale connectivity data was averaged in the subsequent analyses, ensuring a single macroscale data point per tissue sample to match the other samples. All left hemisphere cortical parcels were grouped according to their cortical region category, belonging to multimodal association, unimodal, primary or limbic cortex (Figure 3A). Differences between cortical region categories were assessed by means of Kruskal-Wallis analysis of variance, with post-hoc T-tests for pairwise region category differences (Bonferroni corrected).

**Figure 2.**
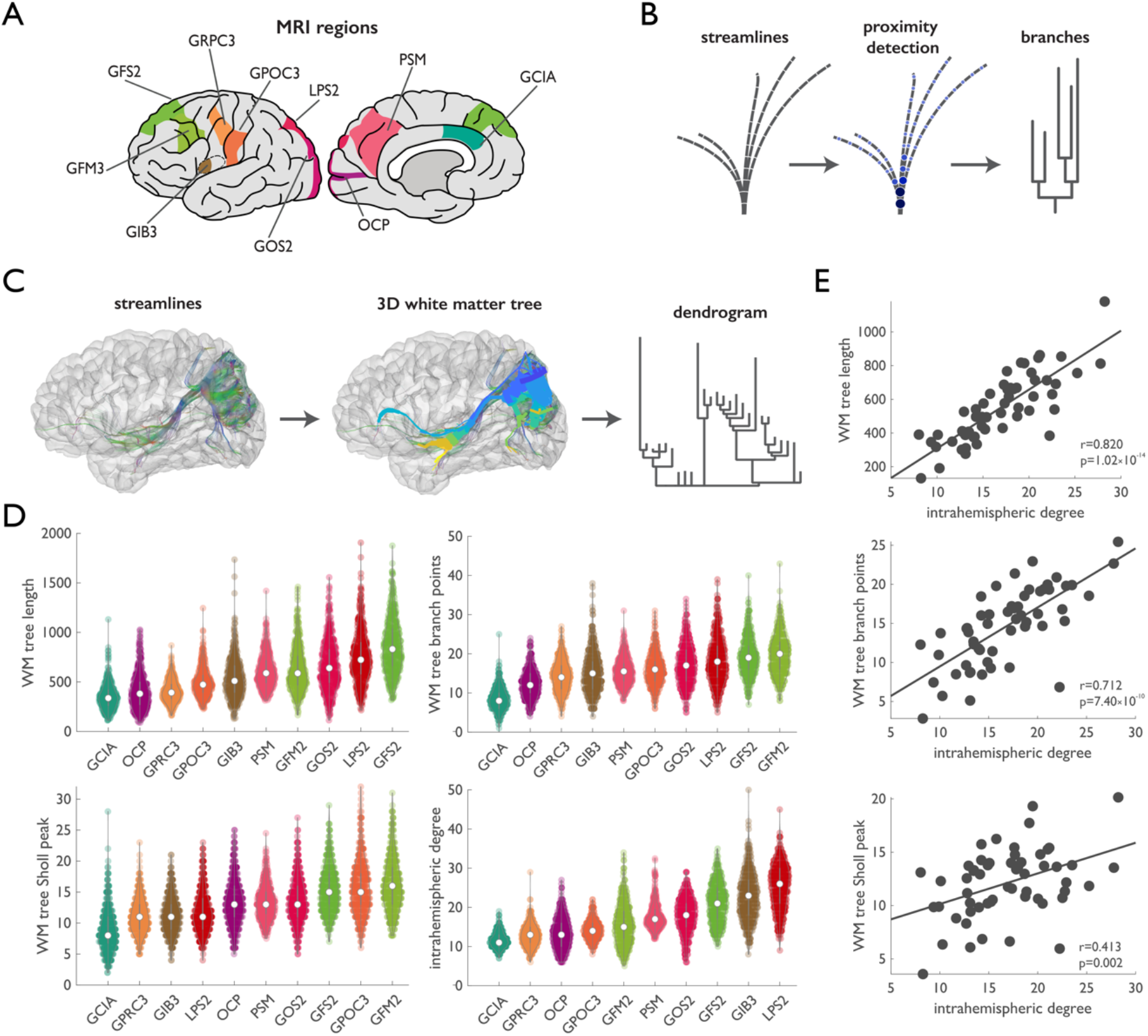
Regional variation in macroscale white matter tree complexity. Panel **A** illustrates the location of the cortical parcels overlapping with the post-mortem sample locations. **B.** Schematic illustration of the reconstruction of a toy white matter tree from a toy set of DWI tractography streamlines, following the trajectory of each streamline in steps of 5 mm (left), and at each step grouping streamlines together when they are within 5 mm distance and projected in the same direction (middle), resulting in a 3D tree suitable for analysis in terms of branching characteristics (right). **C.** From left to right: Visualization of all streamlines touching the superior parietal parcel (*V* in panel A) of a randomly selected HCP subject (ID 107422); the 3D white matter tree resulting from grouping neighboring streamlines; visualization of the branching pattern of the white matter tree. Panel **D** shows regional distributions for all matched-to-neuron samples MRI regions, sorted by region average, depicting white matter tree length (top left), number of branch points (top right), peak Sholl complexity (bottom left) and intrahemispheric degree (bottom right)(Supplemental Figure S3 shows ranked measures of all 57 left hemisphere regions). **E**. Scatterplots showing cross-correlations between degree and the measures of white matter tree complexity, showing a significant correlation with white matter tree length (r=0.820, p=1.02×10-14), branch points (r=0.712, p=7.40×10-10) and peak Sholl complexity (r=0.413, p=0.002)(see Supplemental Figure S4).

**Figure 3.**
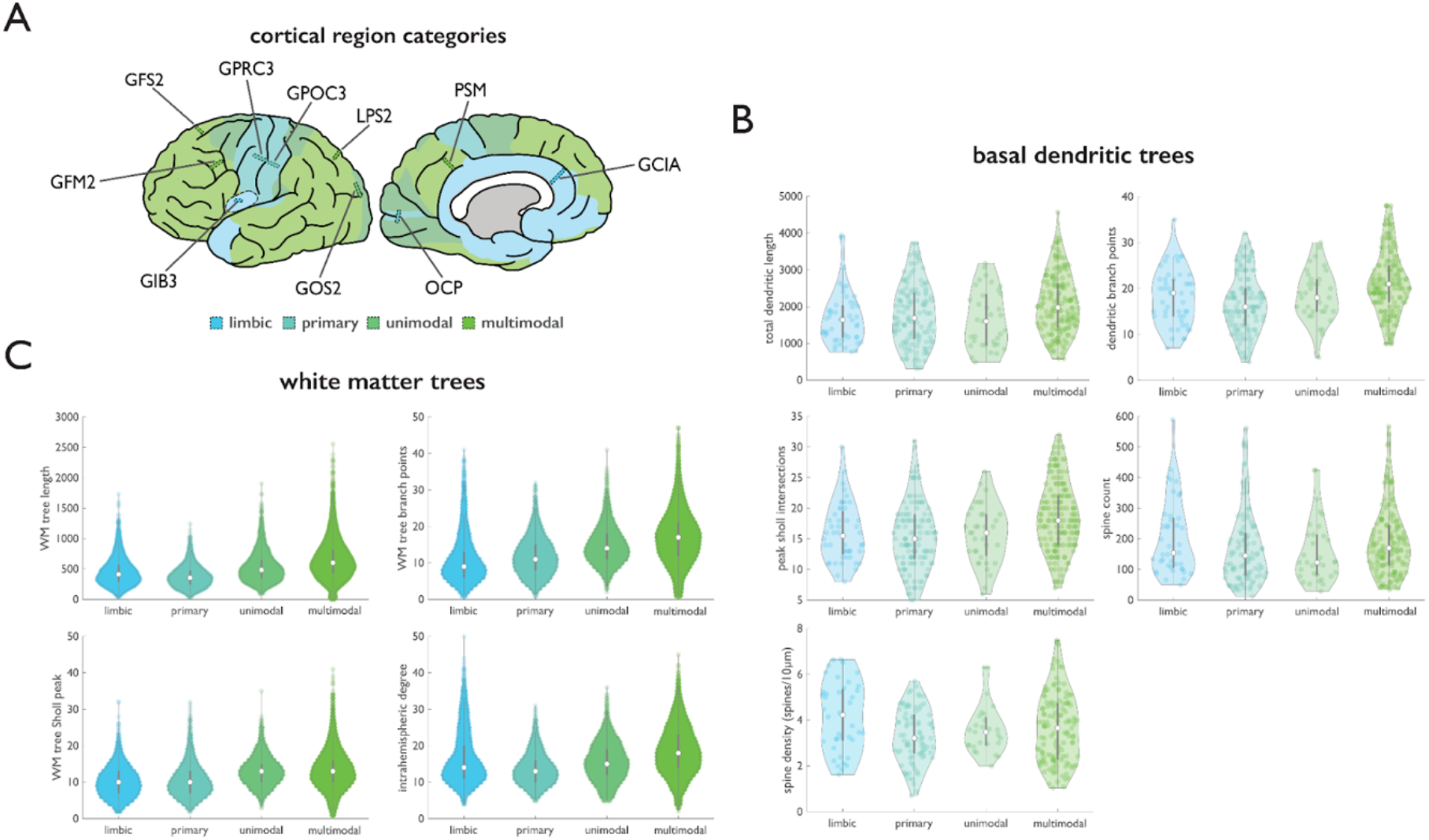
Cross-scale comparison of branching complexity. **A.** Region category assignment into limbic (*GIB3* - anterior insula, *GCIA* - anterior cingulate cortex), primary (*GPRC3* - precentral gyrus, *GPOC3* - postcentral gyrus, *OCP* - occipital pole), unimodal (*GOS2* - superior occipital gyrus) and multimodal association areas (*GFS2* - superior frontal, *GFM2* - middle frontal, *LPS2* - lateral superior parietal, *PSM* - medial superior parietal cortex). **B.** Comparison of microscale basal dendritic tree morphology across cortical region categories, with Kruskal-Wallis non-parametric one-way ANOVA showing significant differences between cortical categories in dendritic tree length (p=0.001), dendritic branch points (p=3.26×10-07), peak Sholl branching complexity (p=5.89×10-06) and spine density (p=0.039), but not spine count. Post-hoc paired comparisons showed that multimodal association areas have neurons with larger dendritic trees than limbic, primary and unimodal cortex (respectively p=0.019, p=0.019, p=0.044), and more branch points and higher peak Sholl complexity than limbic (p=0.046, p=0.018) and primary cortex (p=9.57×10-08, p=7.89×10-06). Spine density was higher in limbic than in primary areas (p=0.023). **C.** Comparison of macroscale white matter tree morphology across cortical region categories, with Kruskal-Wallis non-parametric one-way ANOVA showing significant differences between groups in all four measures (p<0.0001). Post-hoc comparison showed significant differences between all pairs of region categories in white matter tree length (top left panel) and number of white matter branch points (top right panel)(all p<0.0001). Post-hoc comparison in white matter Sholl peak complexity (bottom left) showed limbic and primary regions to have lower Sholl peak complexity than unimodal and multimodal association areas (p<0.0001). Limbic and unimodal areas were observed to have similar degree (bottom right), with significant differences between all other pairs of region categories (p<0.0001).

## Results

### Significant variation in microscale morphology of supragranular pyramidal neurons

Basal dendritic tree length measured on average 1.89×10^03^ μm across reconstructed neurons (std 0.82×10^03^). Dendritic tree length showed significant variation across cortical areas (p=0.001, Kruskal-Wallis analysis of variance), with post-hoc tests confirming significantly larger basal dendritic trees in multimodal association areas compared to primary (p=0.019; Bonferroni corrected), unimodal (p=0.044) and limbic areas (p=0.019)(Figure 3B). Pyramidal neurons with the largest dendritic trees were found in superior frontal gyrus (2.39×10^03^ μm average total dendritic length), and the smallest in the occipital pole (962.1 μm, Figure 1E).

Pyramidal neurons measured on average 19.10 (std 6.82) branch points, showing significant variation across primary, unimodal and multimodal areas (p=3.26×10^−07^). Basal dendritic trees in multimodal association areas were significantly more branched than those in primary (p=9.57×10^−08^) and limbic (p=0.046), but not specifically more than unimodal areas (p=0.282, ns)(Figure 3B). Basal dendrites with highest number of branch points were located in superior frontal gyrus (21.76), and the neurons with the least branched basal dendrites in the occipital pole (12.79)(Figure 1E).

Peak branching complexity (i.e. the location of the peak density of the basal dendritic field of a neuron) was measured to be on average 11.24 (std 9.20). Branching complexity varied consistently across brain areas (p=5.89×10^−06^), with post-hoc testing showing multimodal association areas to have a higher peak branching complexity than areas and primary (p=7.89×10^−06^) and limbic areas (p=0.018), but not unimodal areas (p=0.058)(Figure 3B). Highest peak branching complexity was observed for superior frontal gyrus (17.56 on average), and lowest peak branching complexity was found in anterior cingulate cortex (4.53 on average)(Figure 1E).

Spine count estimated an average 183.1 spines per neuron (quantified on the best visible dendritic branch) (std 115.3), with no significant differences between cortical categories (p=0.058, ns)(Figure 3B). Highest average number of spines was observed in postcentral gyrus (227.6), and lowest in occipital pole (110.3; Figure 1E).

Spine density was on average 3.61 spines/10μm (std 1.39 spines/10μm) across all included neurons. Spine density was observed to be significantly different between region categories (p=0.039), with limbic cortex having higher spine density than primary cortex (p=0.023)(Figure 3B). In line with this observation, of the ten included cortical sample locations, anterior cingulate gyrus was observed to have neurons with on average the highest spine density (4.35 spines/10μm), while precentral gyrus had the lowest (2.96 spines/10μm)(Figure 1E).

### Significant variation in macroscale morphology of white matter structure

We continued by examining the white matter trees for each of the cortical areas (Figure 2B and C), with our reconstructed white matter trees describing the organization of the macroscale connections of that region and providing a template to study regional differences in macroscale branching complexity (Figure 2D, see Supplemental Figure S2 for all n=57 left hemisphere regions). Across all individual white matter trees, average tree length was 545.78 mm (std 279.45), average number of branch points was 14.62 (std 6.72) and peak branching complexity was on average 12.09 intersections (std 4.73). Pooling left hemisphere brain regions of all subjects showed an overall average degree of 16.80 (std 6.26).

We emphasize that white matter tree complexity measures were associated to network degree (i.e. the total number of connections of a region), quantified as white matter tree length (r=0.8204, p=1.02×10^−14^), branch points (r=0.712, p=7.40×10^−10^) and Sholl complexity (r=0.413, p=0.0015), but are by no means identical (Figure 2E and Supplemental Figure S2 and S3). An example is the middle frontal gyrus falling near the middle of the ranking for degree (30th), but scoring much higher on white matter tree complexity (top 9 in branch points and top 4 in Sholl complexity for example).

Next, Kruskal-Wallis non-parametric one-way ANOVA showed significant variation in region categories in all four branching complexity measures (all p<0.0001; Figure 3C). Post-hoc comparison showed significant differences between all pairs of region categories in white matter tree length and number of white matter branch points (all p<0.0001). Post-hoc comparison of the white matter peak complexity showed limbic and primary regions to have lower Sholl peak complexity than unimodal and association areas (p<0.0001). Limbic and unimodal areas were observed to have similar degree, with significant differences between all other pairs of region categories (p<0.0001).

We ranked the ten matched-to-histology macroscale cortical regions (Table 2, Figure 2A) in terms of the branching complexity of their associated white matter trees (Figure 2D). Rostral middle frontal gyrus displayed on average the most branched (19.79, std 5.04) white matter trees with the highest Sholl complexity (16.24, std 4.26), while superior frontal gyrus had the longest white matter trees (862.65 mm, std 276.60 mm) and superior parietal cortex the highest macroscale degree (25.25, std 5.76). Smallest white matter tree structures were observed in anterior cingulate (length 349.31 mm, std 128.14 mm; branch points 8.48, std 2.90; Sholl complexity 8.81, std 3.47; degree 11.66, std 1.95), followed by precentral gyrus (second lowest Sholl complexity and degree; respectively 11.16 (std 2.80) and 12.80 (std 2.77)) and primary visual cortex (second lowest total tree length and branch points; 406.41 mm (std 180.55 mm) and 12.33 (std 3.68))(Figure 2D).

### Cross-scale analysis shows micro-macro correlation in connectivity

Significant cross-scale associations were found in the regional patterns of branching complexity, with the number of branch points showing strongest correspondence across scales (r=0.797, p=0.006; Figure 4F). Number of connections as quantified by microscale spine count and macroscale degree; (r=0.677, p=0.032; Figure 4H), total tree length (r=0.664, p=0.036; Supplemental Figure S4) and peak Sholl branching complexity (r=0.724, p=0.018; Figure 5) also showed significant multi-scale association. Number of branch points showed particularly a strong association across scales, also with other non-equivalent measures of branching complexity (Table 3). Branch points of pyramidal neuron basal dendrites was associated with macroscale regional degree (r=0.677, p=0.032), white matter branch points (as reported above, Figure 4E-F) and white matter tree length (r=0.798, p=0.006). In turn, white matter branch points showed strong association with microscale spine count (r=0.728, p=0.017), dendritic branch points (as reported above), dendritic length (r=0.762, p=0.010) and dendritic peak Sholl branching complexity (r=0.766, p=0.009), but not spine density (see Supplemental Results).

**Figure 4.**
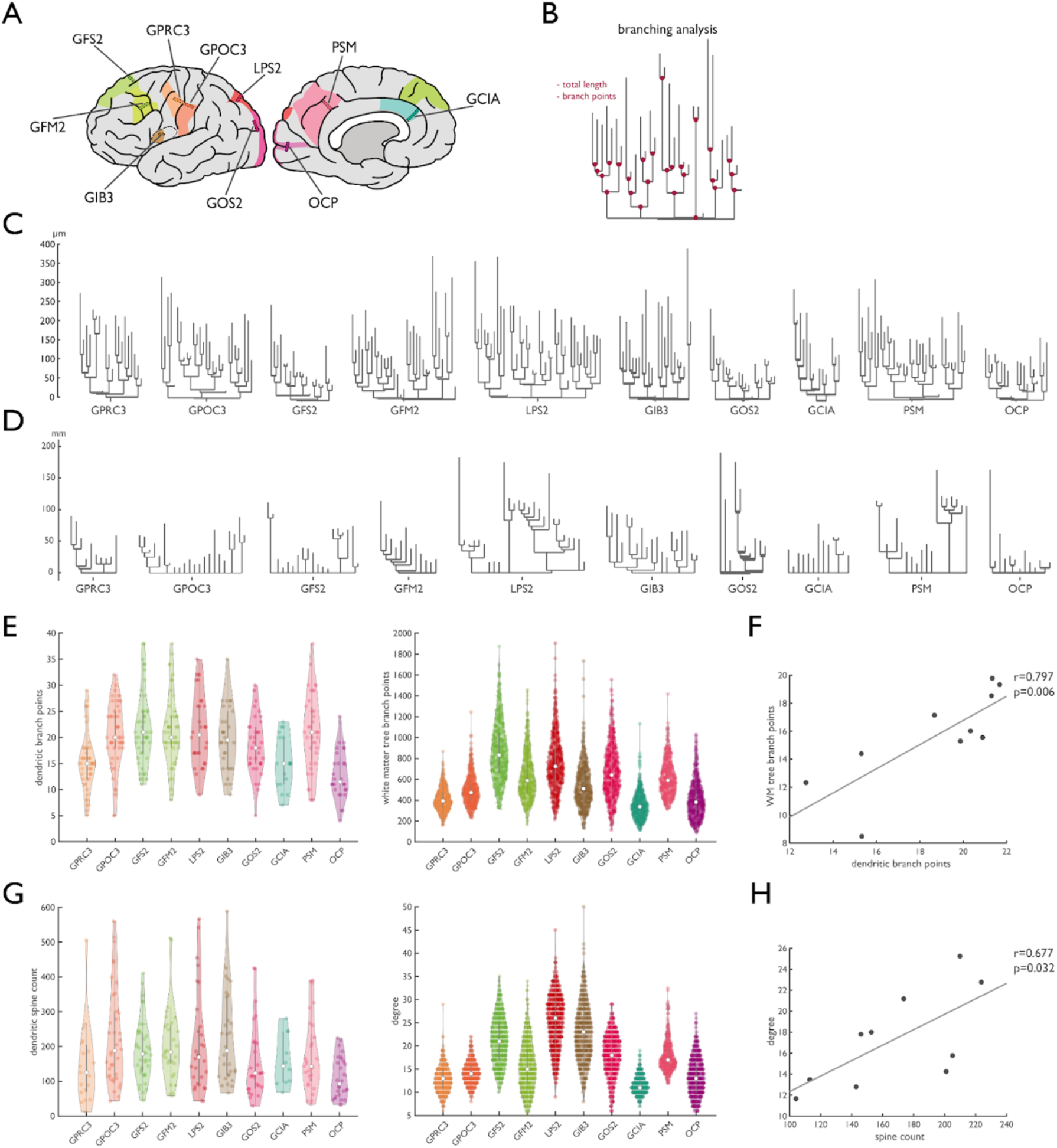
Similar regional dendritic and white matter branching. Panel **A** shows a schematic illustration of the locations of the matching donor sample locations (small rectangles) and FreeSurfer parcels (colored patches) for the ten included brain regions. Panel **B** illustrates two measures of neural branching: the total length and the number of branch points. **C** shows dendrograms of exemplary neurons for all regions depicted in A (same neurons as in Figure 1C), exemplary dendrograms for white matter trees of those same 10 brain regions (of HCP participant 100307) are depicted in panel **D**. Panel **E** shows the regional variation in number of branch points in the reconstructed neurons (left) and in the macro-scale white matter trees (right). The regional number of branch points shows considerable multi-scale overlap (r=0.797, p=0.006) (**F**). Panel **G** shows the regional distribution of dendritic spine count (left) and macroscale degree (right), with the cross-scale association (r=0.676, p=0.032) depicted in **H**.

**Figure 5.**
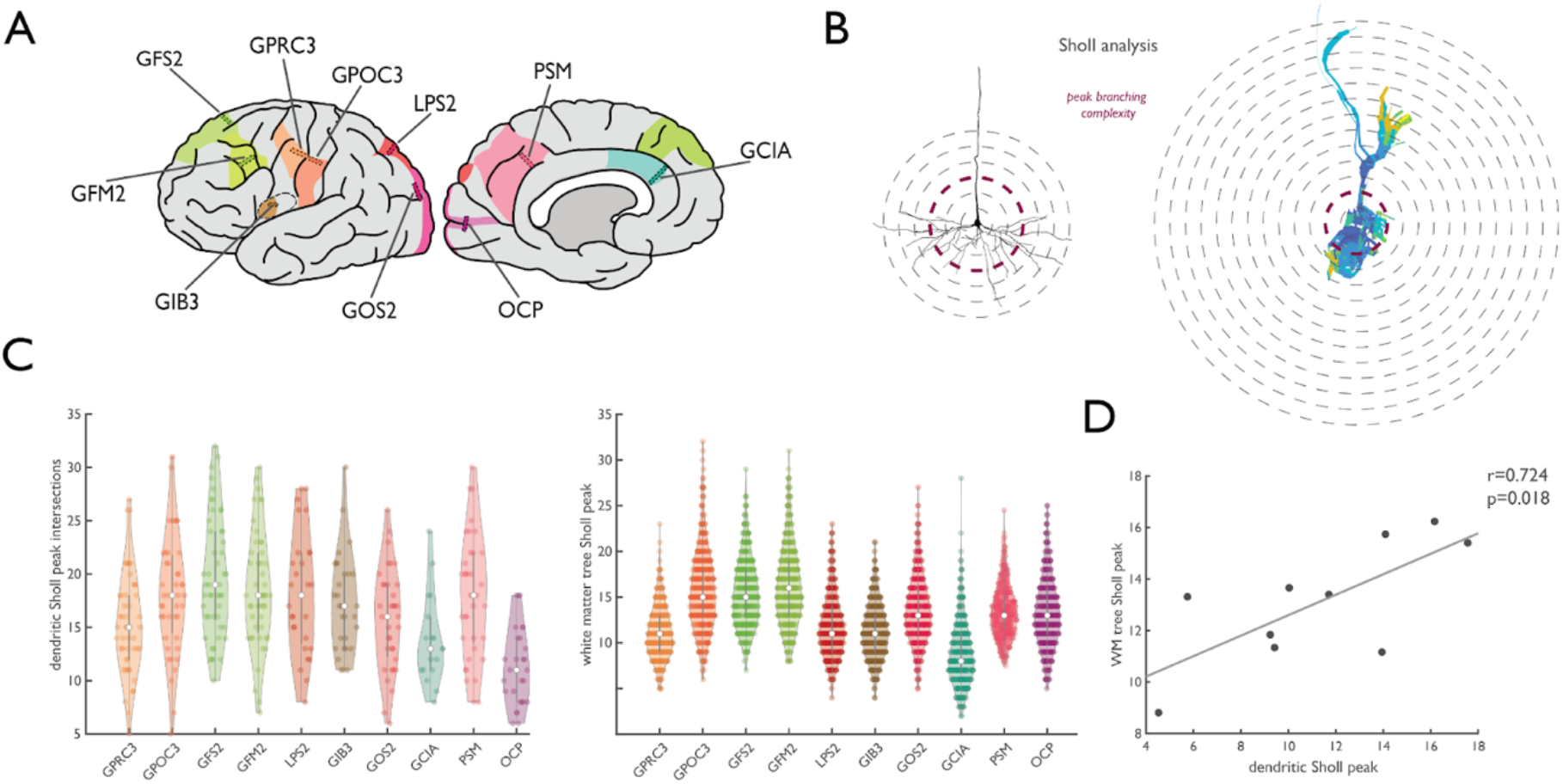
Similar regional dendritic and white matter Sholl complexity. Panel **A** shows a schematic illustration of the locations of the matching donor sample locations (small rectangles) and FreeSurfer parcels (colored patches) for the ten included brain regions. **B**. Schematic depiction of Sholl analysis on neuron basal dendrite (left) and white matter tree branching (right). Panel **C** shows the regional variation in peak Sholl intersections in the reconstructed neuron basal dendrites (left) and in the macro-scale white matter trees (right), with considerable multi-scale overlap (r=0.724, p=0.018) (**D**).

**Table 3.** Cross-scale associations between neuronal and white matter branching structure, listing Pearson’s *r* and *p*-value.

|  | degree | WM tree<br>branch points | WM tree<br>length | WM tree<br>Sholl peak |
| --- | --- | --- | --- | --- |
| spine count | **0.676 (0.032)** | **0.727 (0.017)** | 0.521 (0.122) | 0.409 (0.241) |
| dendritic branch points | **0.637 (0.032)** | **0.797 (0.006)** | **0.798 (0.006)** | 0.519 (0.124) |
| dendritic length ($\mu\text{m}$) | 0.464 (0.171) | **0.762 (0.010)** | **0.664 (0.036)** | 0.466 (0.174) |
| dendritic Sholl peak | 0.162 (0.654) | **0.766 (0.010)** | 0.549 (0.100) | **0.724 (0.018)** |
| spine density (spines/10$\mu\text{m}$) | 0.152 (0.675) | -0.225 (0.532) | -0.024 (0.948) | -0.306 (0.390) |

## Discussion

Our findings point to similar characteristics of brain connectivity across scales. White matter macroscale wiring patterns display similar connectivity characteristics as the neuronal microscale. Well-known aspects of neuronal organization that impact local information processing such as the length of dendritic wiring, branch points and branching complexity, were found to scale in a similar manner across macroscale areas.

Our observed cross-scale relationship in connectivity complexity is supported by empirical and computational modeling studies at both the cellular and whole brain level. Economy of brain wiring has been proposed to play an important guiding role in shaping brain connectivity. Both cellular and neuroimaging studies have separately shown brain connectivity to display small-world, modular and scale-free properties, demonstrating a trade-off between a minimization of wiring cost and processing efficiency (Bullmore and Sporns, 2012; van den Heuvel et al., 2012). Wiring economy in microscale pyramidal neuron morphology balances efficient signal integration across the neuron with signal segregation through local processing in dendritic segments and branches (Branco and Häusser, 2010). Dendritic tree length influences a neuron’s response to incoming information (Amatrudo et al., 2012), with shorter dendritic trees more suited for rapid coincidence detection, while larger, more branched dendritic trees increase a cell’s capacity for (slower) temporal integration (Amatrudo et al., 2012; Papoutsi et al., 2017). At the system level, similar organizational principles are observed in cell-to-cell networks (Towlson et al., 2013; Shih et al., 2015), hippocampal networks (Bonifazi et al., 2009) and in simulated cortical micro-connectomes (Gal et al., 2017). In support of our findings of association regions to show both higher neuronal complexity and more and more complex macroscale connectivity, higher-order association areas are generally characterized by a highly diverse structural and functional connection profile, ideal for a central role in integrative information processing in the brain (Felleman and Van Essen, 1991; van den Heuvel and Sporns, 2013; Zhang et al., 2019). Cortical regions involved in primary sensory processing have a more focused connectivity profile, ensuring initial more segregated processing of sensory inputs (Felleman and Van Essen, 1991), which again fits with our current observations of relatively less complex microstructural and macrostructural connectivity profiles of primary regions, indicating that indeed micro and macroscale branching complexity go hand in hand.

Studies modeling macroscale region-to-region connectivity of the brain have suggested that the connections of a cortical region are shaped by multiple factors. One guiding factor is the spatial distance between brain regions, with regions close to each other being more likely and more stronger connected than more distant regions (Beul et al., 2017; Goulas et al., 2019). The likelihood of being connected further increases when regions are of similar cortical type (Barbas and Rempel-Clower, 1997). A second proposed mechanism involves physical processes such as tension-based morphogenesis, proposing that the compact morphological structure of white matter bundles is the result of the shaping force of physical tension along the length of developing axons and dendrites, reaching across scales of organization to bring connected regions closer together (Essen, 1997). Alternatively, the three-dimensional structure of brain wiring has been described as a spatially continuous grid, established along the three primary chemotactic gradients of early embryogenesis and deformed by cerebral expansion in later development (Van Wedeen et al., 2012).

A similar continuity in the organization of brain connectivity is observed on the microscale. Primate studies have shown that the size of a pyramidal neuron’s (basal) dendritic tree and the reach of its local, intra-areal axonal projections are strongly related (e.g. (Lund et al., 1993)) and increase along the cortical hierarchy (Elston, 2003b). The extent of a neuron’s local and long-range connectivity go hand in hand as well, and together show an increase in breadth of neuronal connectivity from primary to association areas (Amir et al., 1993). These findings, together with our observation of consistent branching complexity across the micro and macroscale of the connectome, support the notion of a continuous, consistently organized wiring structure of the human brain across scales.

A number of practical limitations should be considered. The majority of brain donors are of relatively advanced age (mean 78.8, range 55-98 years). Supragranular pyramidal neurons tend to be relatively unaffected by ageing compared to those in infragranular layers (de Brabander et al., 1998). The overall regional pattern of larger dendrites and more spines in higher order areas, which is the focus of the current study, remains rather robust (Jacobs et al., 1997; Jacobs et al., 2001). It should be noted that the three oldest tissue donors displayed at least some degree of neuropathological changes (see also Table 1). The oldest two tissue donors were later diagnosed with respectively early stage and advanced Alzheimer’s disorder (AD) based on these neuropathological hallmarks. Excluding these individuals did not change our findings. Validation analyses showed that although overall dendritic complexity in the older donors was lower than in the younger donors (Supplemental Figure S6), this had no effect on the presented results (Supplemental Figure S7).

Basal dendritic branches of supragranular pyramidal neurons were reconstructed in sections of Golgi-Cox stained tissue samples (Glaser and Van der Loos, 1981). This procedure is believed to stain a random subset of neurons, leaving the majority of neurons unstained and translucent, thus enabling the visual tracing and 3D reconstruction of neuron morphology. The mechanism by which some neurons are stained while others are not remains elusive (see for discussion e.g. (Ramon-Moliner, 1970; Buell, 1982; Swaab and Uylings, 1987)). The Golgi-Cox stained tissue allowed for reconstruction of primarily lateral basal dendrites projecting more or less horizontally from the soma and thus results in a partial reconstruction of the dendritic tree.

We further note that diffusion-weighted imaging is known to provide only an indirect reconstruction of anatomical connectivity compared to more direct methods such as histological tract-tracing (Donahue et al., 2016). Fiber orientations modeled using diffusion weighted imaging have however been shown to show correspondence with histological myelinated fiber orientation in the human (Seehaus et al., 2015) and primate brain (Kaufman et al., 2005) and μCT axon orientation in mouse brain (Foxley et al., 2020) and metrics of DWI derived metrics of connectivity strength tend to show overlap with metrics derived from more gold-standard tracer-based connectivity measurements in for example the macaque cortex (van den Heuvel et al., 2015; Delettre et al., 2019).

Our analyses were performed using a combination of *in vivo* high-resolution diffusion-weighted MRI datasets of relatively young (aged 20-35 years) participants of the Human Connectome Project (Van Essen et al., 2013) and *ex-vivo* cortical samples of older donor brains (aged 55-98 years). Because of this cross-cohort study design, our observations should be taken as a general indication of consistency in organization between neuronal microscale and region-to-region macroscale wiring. As with any association-based study, the question arises of how the observed shared micro-macro organization may have come to be, what the causal direction of the association is, or whether perhaps a third underlying factor guides both. Future combined neuronal and white matter reconstruction within the same individuals would be of high interest to further substantiate the made observations.

Our findings suggest the existence of multi-scale principles of connectivity organization in the human brain. A first question of high interest is how such observed shared principles might come to be, both from a perspective of evolution and of brain development. Future comparisons incorporating the interplay of connectivity organization with other neuronal and cell types are also of great interest. Other cell types (e.g. inhibitory neurons, astrocytes) have a crucial role in regulating activity within a brain region and play an important role in the branching and spiking behavior of pyramidal neurons (Bonifazi et al., 2009; Kanner et al., 2018; Hafizi et al., 2021; Kajiwara et al., 2021), as well as in disease (e.g. (Marín, 2012; Dossi et al., 2018)).

Cross-scale investigations of connectivity may help to get a better understanding of disease processes that affect brain connectivity. Many neurological and psychiatric conditions are reported to show changes in brain connectivity at both the cellular and macroscale level, for example in Alzheimer’s disease (e.g. (Braak and Braak, 1996; Uylings and de Brabander, 2002; He et al., 2008; Stam et al., 2009)) and schizophrenia (e.g. (Garey et al., 1998; Glantz and Lewis, 2000; Whitfield-Gabrieli et al., 2009; van den Heuvel et al., 2010)). Studies examining neurobiological correlates of brain disorders have shown that regional patient-control differences in neuroimaging-based markers of cortical morphometry and connectivity coincide with microscale differences in those areas, for example in Alzheimer’s disease (Buckner et al., 2009; Prescott et al., 2014) and schizophrenia (van den Heuvel et al., 2016). White matter tree complexity is a novel measure of interest to study patient-control differences in brain wiring. In particular neurodevelopmental disorders could benefit from examining potential developmental perturbations of the branching structure of white matter bundles, as suggested for changes in neural connectivity at the microscale level (Selemon and Goldman-Rakic, 1999; Zikopoulos and Barbas, 2013; Petanjek et al., 2019).

## Acknowledgments

We thank Dr. Corbert van Eden for his indispensable practical guidance in the lab, for sharing his knowledge on the staining protocols and for his insightful comments that helped make this research possible. M.P.v.d.H. is an MQ fellow and was supported by an Innovational Research Incentives Scheme VIDI grant (Grant no. VIDI-452-16-015) of the Netherlands Organisation of Scientific Research (Nederlandse Organisatie voor Wetenschappelijk Onderzoek). M.P.v.d.H. was supported by an ALW Open grant (Grant no. ALWOP.179) of the Netherlands Organisation of Scientific Research. Human data were provided by the Human Connectome Project, WU-Minn Consortium (Principal Investigators: David Van Essen and Kamil Ugurbil; 1U54MH091657) funded by the 16 NIH Institutes and Centers that support the NIH Blueprint for Neuroscience Research; and by the McDonnell Center for Systems Neuroscience at Washington University.

