## Supplemental materials for "Common micro- and macroscale principles of connectivity in the human brain"

Lianne H. Scholtens, Boelelaan 1085, W&N B-636, 1081 HV, Amsterdam, The Netherlands

**Supplemental Methods**

Golgi-Cox staining protocol

*Tissue dissection.* Tissue samples were dissected in 0.5 cm thin blocks perpendicular to the gyral surface spanning the width of the gyrus of interest, while avoiding any unnecessary pressure on the tissue to minimize staining artefacts later in the Golgi-Cox staining process. For donors 3-5, a photo was taken of the location of the dissection and the brain tissue was moved directly into light-proof jars containing Golgi-Cox solution, for simultaneous fixation and impregnation (Van der Loos, 1959).

*Golgi-Cox impregnation.* After 48 hours of impregnation, Golgi-Cox solution was replaced to reduce tissue precipitate. After 2.5 to 3 weeks of Golgi-Cox impregnation, the quality and progression of the Golgi-Cox impregnation was assessed. Criteria for Golgi-Cox impregnation quality assessment were 1) complete impregnation of the dendritic branches (including the tips of both basal and apical dendrites) of cells throughout the section; 2) clearly visible dendritic spines on all most distal branches; while 3) avoiding over-impregnation of the tissue. When the impregnation was judged to be complete, tissue blocks were removed from the Golgi-Cox solution, rinsed briefly in tap water and moved to the next tissue processing step. In the case that the neuronal branches were not yet sufficiently impregnated, tissue blocks were kept in the Golgi-Cox solution for an additional 1 to 2 days, after which the impregnation was reassessed.

*Dehydration and celloidin embedding.* Impregnated tissue samples were dehydrated over the course of 2 days in a graded ethanol series (ethanol 70%-96%-100%), followed by 24 hours in a mixture of 100% ethanol – (di)Ethyl ether (1:2). After dehydration, samples were placed in a solution of 6% celloidin in 100% ethanol – (di)Ethyl ether (1:2) for 5 days, followed by 7 days in a 10% celloidin solution. For embedding, each individual sample was placed in a paper box made to size. Boxes were made of thick uncoated paper to allow for the penetration of chloroform and ethanol in the subsequent steps. Each box was filled with a thin base layer of 10% celloidin, which was allowed some time to set in order to form a thin membrane before carefully placing a tissue sample on top. Samples were oriented to later facilitate the preferred plane of cutting in the sectioning step, the box was further filled with 10% celloidin to cover the sample, which was allowed to set for 30 minutes. Next, boxes were placed in a glass desiccator with weights placed on top and submerged in chloroform. After hardening in chloroform for 24 hours, boxes were moved to ethanol 70% for storage.

*Sectioning.* Before sectioning, tissue blocks were removed from their paper box, mounted onto a block holder using 10% celloidin and placed in ethanol 70% overnight until fixed. Tissue blocks were sectioned perpendicular to the cortical surface, and parallel to the apical dendrites of the pyramidal cells. The tissue was sectioned into 180 µm thick sections using a sledge microtome (Leica / Reichert-Jung Polycut S) and transferred to ethanol 70% in individual compartments of a Teflon disk for the free-floating development and dehydration process. The sections were rinsed in demineralized water, after which the staining was developed using a 16% ammonium solution ($NH_4OH$; 15 minutes), followed by rinsing and a 1% sodium thiosulfate bath ($Na_2S_2O_3$; 7 minutes). Next, the sections were again thoroughly rinsed, after which the sections for Nissl counter staining were moved to the cresyl violet solution, while the rest was moved through 5 minutes each of ethanol 70%, ethanol 90% and butanol for dehydration. Finally, the sections were placed in Histoclear (National Diagnostics, Atlanta USA) for 5 minutes, after which they were mounted in Histomount (National Diagnostics, Atlanta USA) on microscopic slides and coverslipped. Weights were placed on the coverslips in order to smooth out the sections, and the slides were placed in the dark at 4°C to dry. After drying, the weights were removed, the slides cleaned of excess mounting medium and stored horizontally in the dark at 4°C.

*Nissl counterstaining of non-impregnated neurons.* In every sixth section, non-impregnated neurons were counterstained using a solution of 2.5% cresyl violet, in order to provide assessment of the cortical layering in the sample. The sections were submerged in the cresyl violet solution for 10 minutes, at which point the tissue was stained completely purple. After rinsing off the excess dye in demineralized water, the sections were quickly rinsed in 70% ethanol until all background staining had been cleared, after which the sections were dehydrated and further processed according to the main protocol (see above paragraph).

**Supplemental Results**

*Spine density micro-macro correlation*

Cortical variation in spine density was not found to show a clear association with macroscale wiring organization when all ten cortical regions. Anterior cingulate and anterior insular regions were however observed as relative outliers compared to the other samples, with respectively highest and second highest spine density, but relatively ranked low in terms of cortico-cortical macroscale connections. Post-hoc analysis excluding these limbic regions from the spine density x macroscale wiring comparison revealed a significant association between spine density and white matter tree branch points ($r=0.792$ $p=0.019$; Figure S5), with possible trend-level associations with white matter tree length ($r=0.673$, $p=0.068$) and peak Sholl complexity ($r=0.669$, $p=0.069$).

### Supplemental Figures

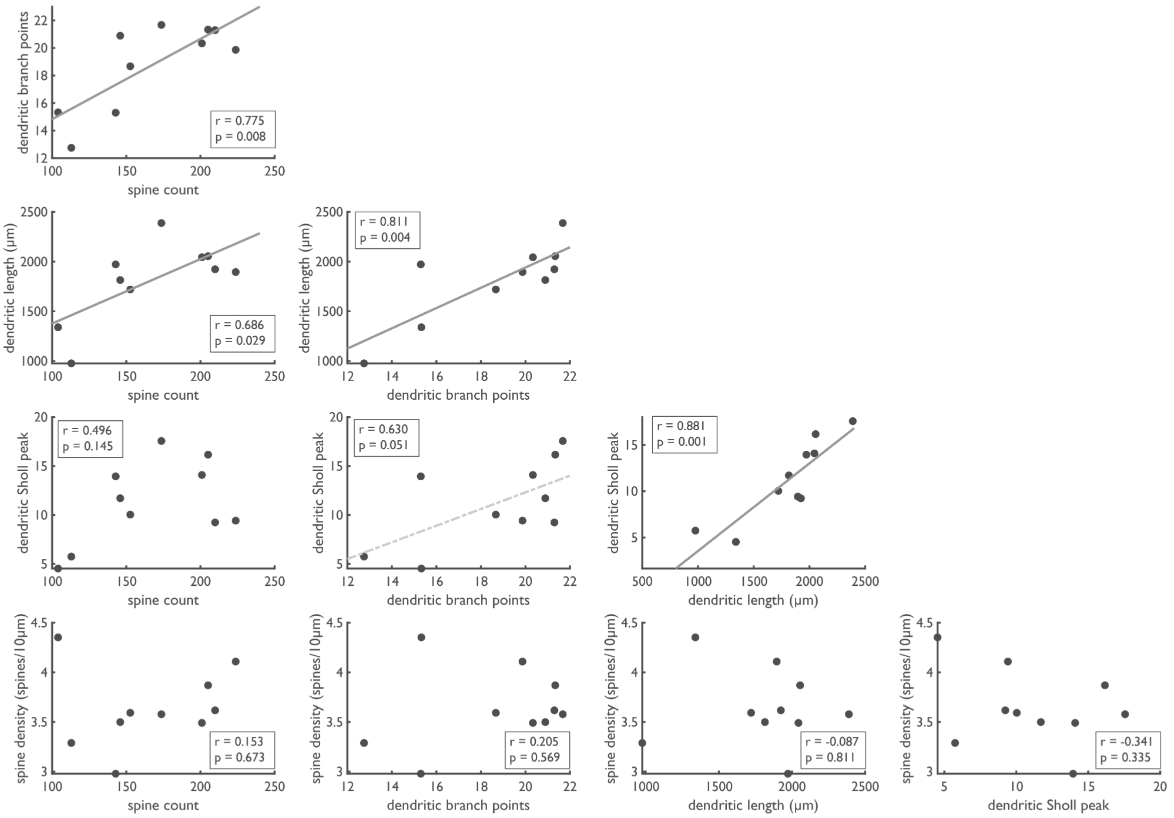


**Supplemental Figure S1. Associations between microscale measures of neural wiring complexity.** Cross-correlation shows associations between the majority dendritic branching complexity measures, with the notable exception of Sholl peak complexity and spine density.

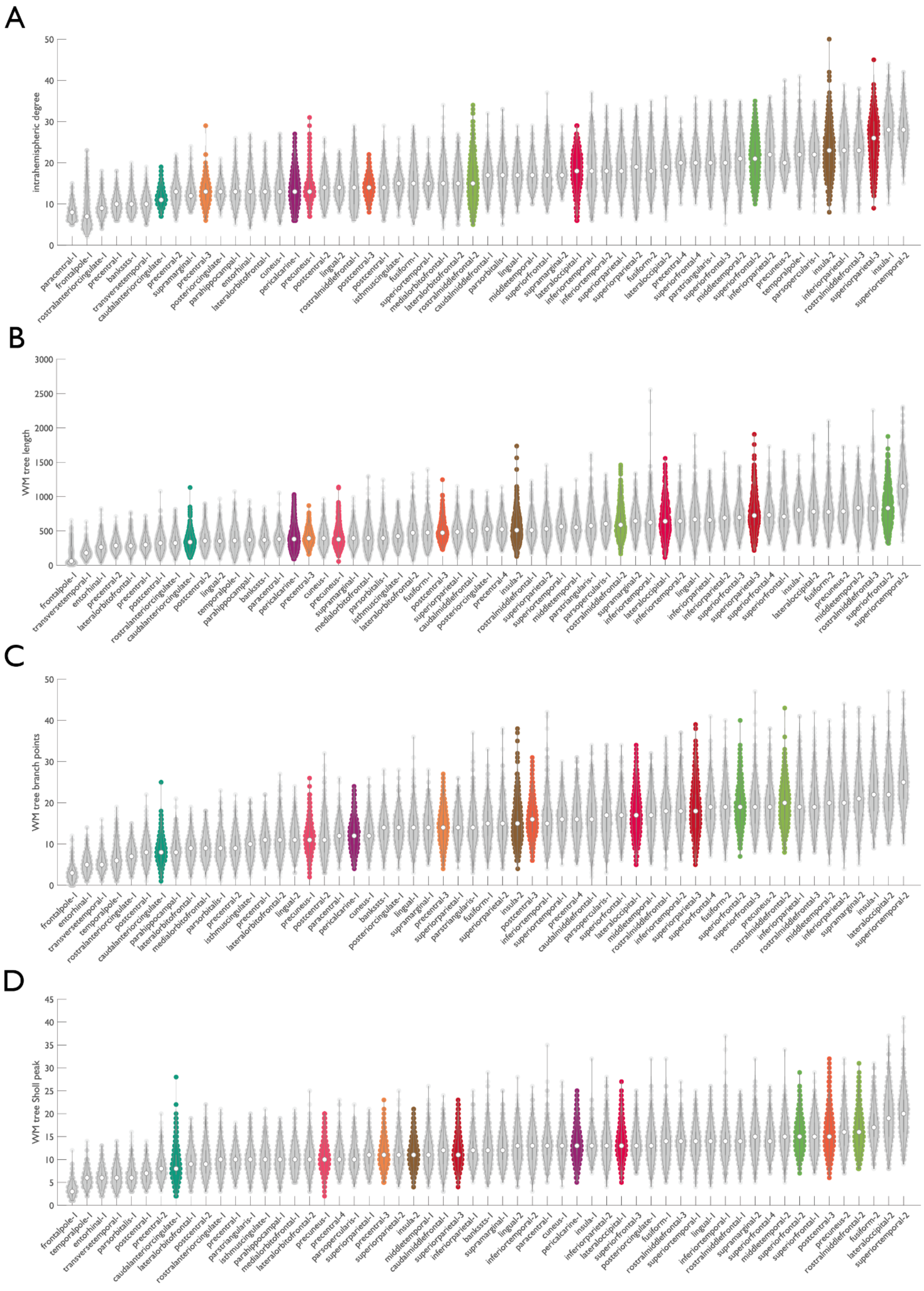


**Supplemental Figure S2. Ranking of macroscale Cammoun-114 parcels by neural branching complexity.** Within each violin, individual points represent HCP datasets, with the dot indicating the median value. Colored violins highlight regions included in the multiscale branching analysis.

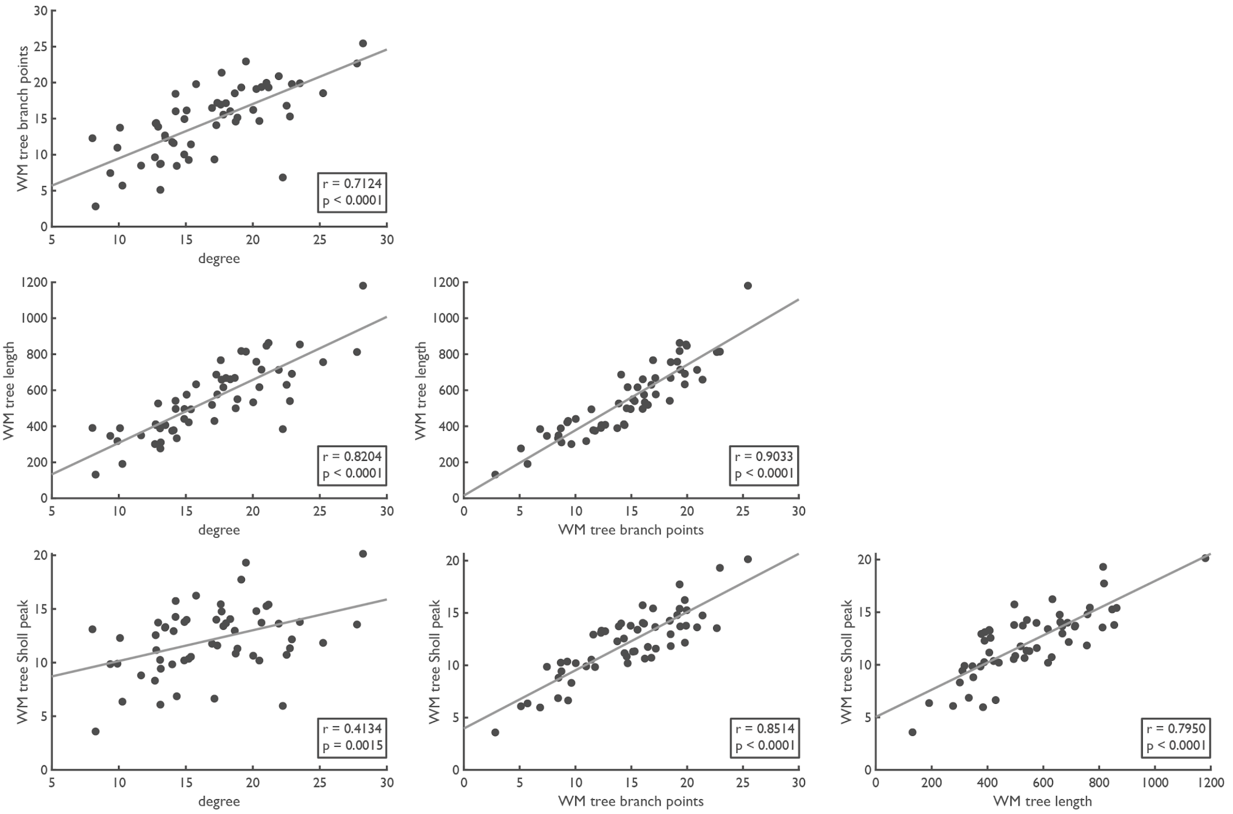


**Supplemental Figure S3. Associations between macroscale measures of neural wiring complexity.** Cross-correlation shows strong correlations between the macroscale connectivity and branching complexity measures (all p<0.0001, except white matter Sholl peak complexity x degree (p=0.0015)).

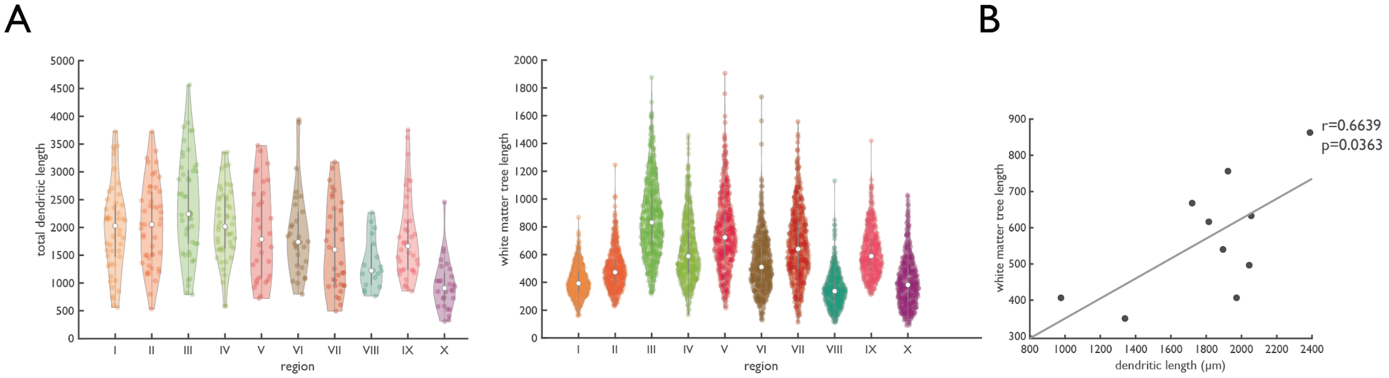


**Supplemental Figure S4. Similar dendritic and white matter tree length across cortical regions.** Panel **A** shows the regional distribution of total dendritic length (left) and macroscale white matter tree length (right). The scatter plot in panel **B** shows the cross-scale association, with dendritic length on the x-axis and white matter tree length on the y-axis (r=0.6639, p=0.0363).

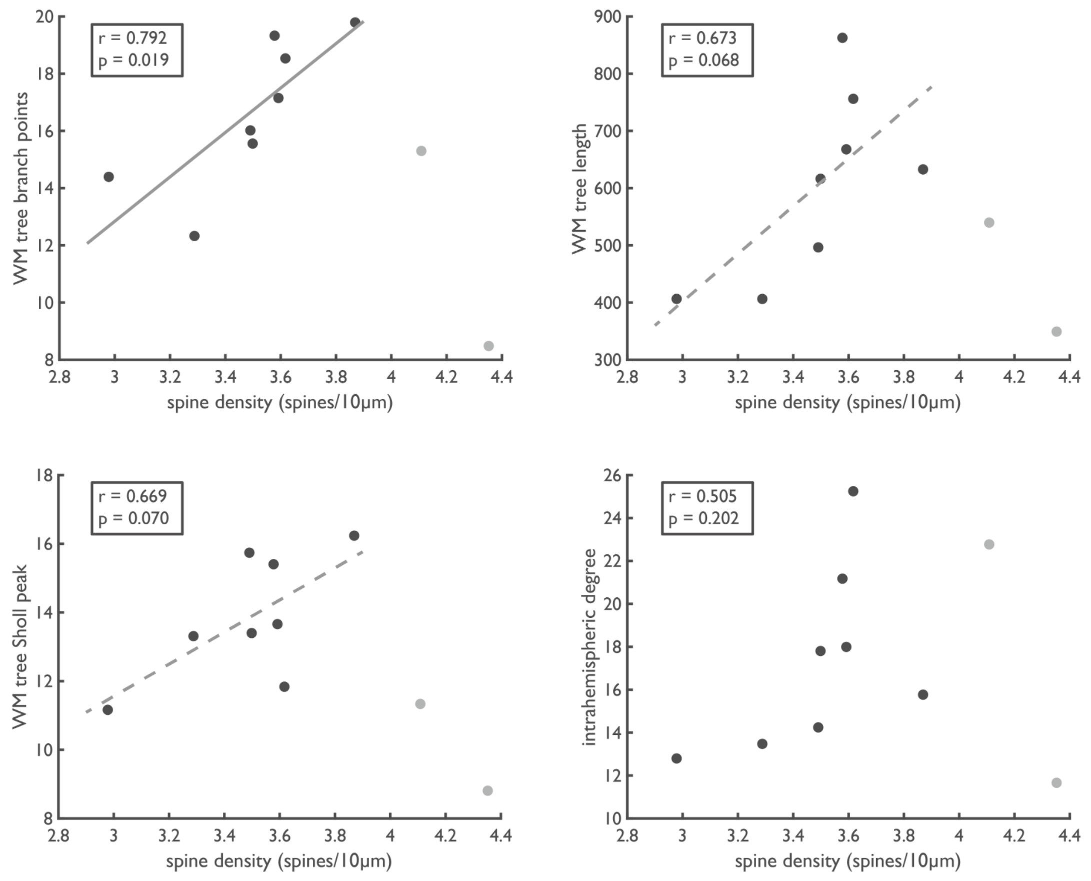


**Supplemental Figure S5. Spine density and macroscale complexity.** Figure shows scatter plots of the micro-macro association between basal spine density (spines/10μm) and the four measures of macroscale branching complexity. Limbic areas anterior cingulate cortex and anterior insula were observed to be relative outliers, ranking very high spine density but low on the other neuronal complexity measures (see Figure 1E). Post-hoc analysis excluding these points from the regional cross-scale comparison showed an association between spine density and the number of white matter tree branch points (r=0.792, p=0.019; top left panel), with trend-level associations with white matter tree length (r=0.673, p=0.068; top right) and white matter peak Sholl complexity (r=0.668, p=0.070; bottom left), but not intrahemispheric degree (r=0.505, p=0.202; bottom right).

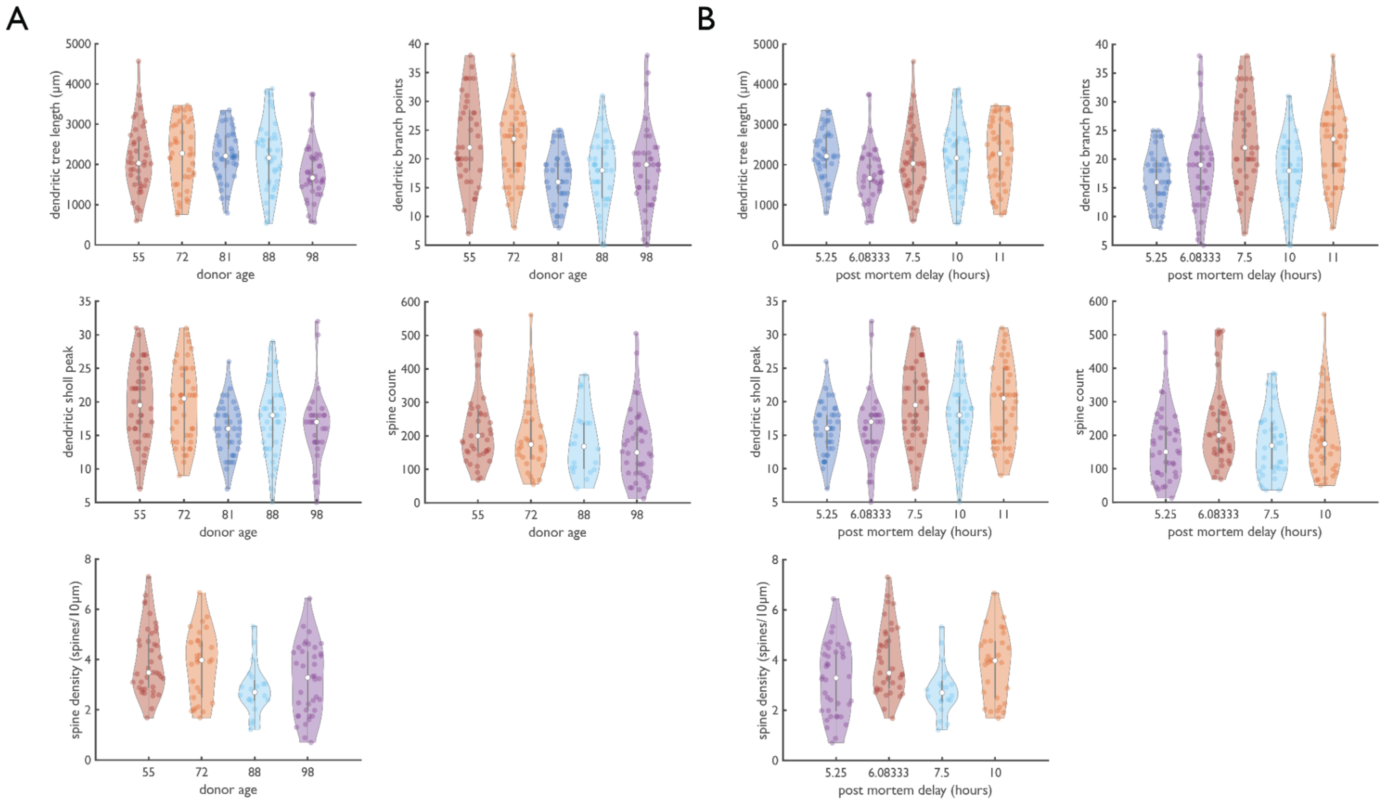


**Supplemental Figure S6. Effect of donor age and post mortem delay.** Figure shows the individual neuron data aggregated over the four regions for which data of all donors was available: precentral gyrus, postcentral gyrus, superior frontal gyrus and middle frontal gyrus, sorted by donor age (panel **A**) and by post mortem delay (panel **B**). Kruskal-Wallis non-parametric one-way ANOVA shows significant differences between the donors in dendritic tree length ($p = 5.2\times10^{-5}$, with the oldest donor having shorter basal dendrites), dendritic branch points ($p = 0.024$), peak Sholl complexity ($p = 0.0052$; older donors (aged 80+) tend to have neurons with lower complexity) and spine density ($p = 0.0058$), but not spine count ($p = 0.081$). There appear to be no systematic differences in neuron branching complexity related to post mortem delay.

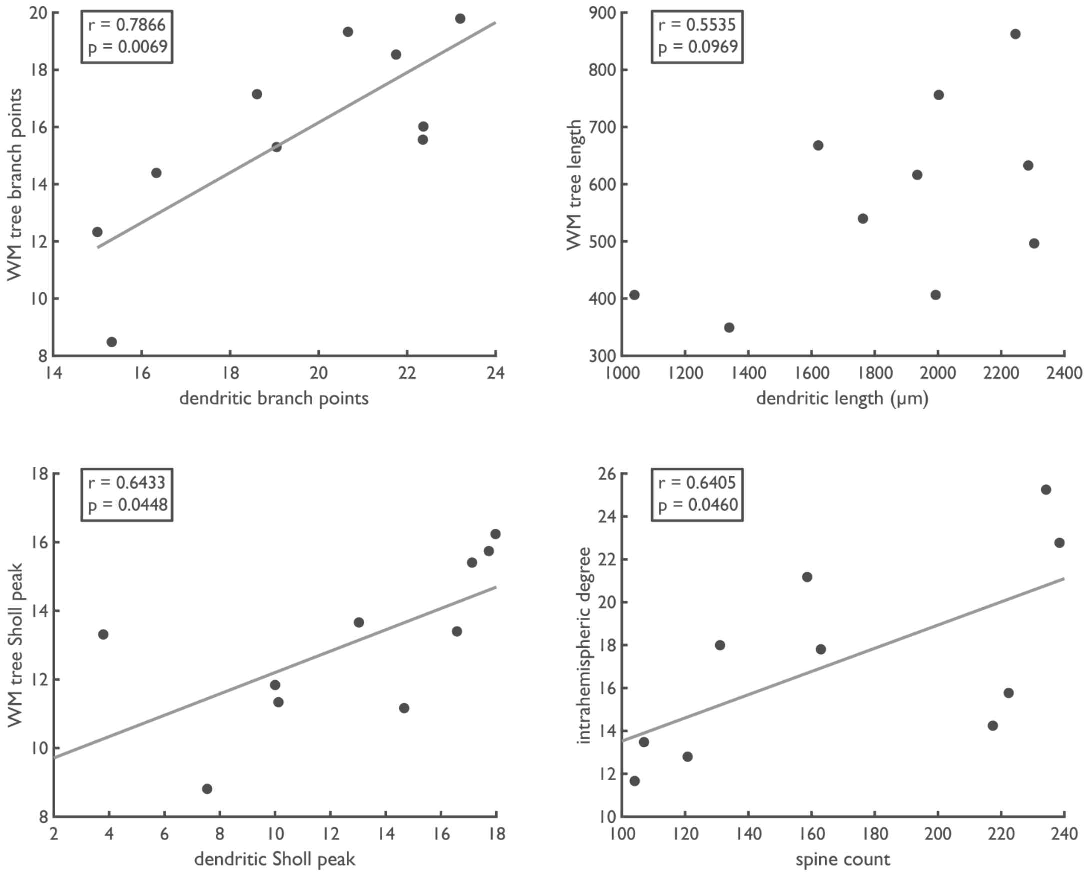


**Supplemental Figure S7. Analysis without Alzheimer's disease diagnosed donors.** Figure shows the cross-scale association between neural and white matter branching complexity, performed leaving out the donors diagnosed with Alzheimer's disease (donor 2 and 3, also the oldest included donors at 88 and 98 years old). Leaving out these donors yielded results highly similar to the main analysis, though without significant association between dendritic- and white matter tree length.
